# Type-2-diabetes Alters CSF but not Plasma Metabolomic and AD Risk Profiles in Vervet Monkeys

**DOI:** 10.1101/665117

**Authors:** Kylie Kavanagh, Stephen M. Day, Morgan C. Pait, William R. Mortiz, Christopher B. Newgard, Olga Ilkayeva, Donald A. Mcclain, Shannon L. Macauley

## Abstract

Epidemiological studies suggest that individuals with type 2 diabetes (T2D) have a 2-4 fold increased risk for developing Alzheimer’s disease (AD), however the exact mechanisms linking the two disease is unknown. In both conditions, the majority of pathophysiological changes (including glucose and insulin dysregulation, insulin resistance, and AD-related changes in Aβ and tau) occur decades before the onset of clinical symptoms and diagnosis. In this study, we investigated the relationship between metabolic biomarkers associated with T2D and AD-related pathology, including Aβ levels, from cerebrospinal fluid (CSF) and fasting plasma of healthy, prediabetic (PreD), and T2D vervet monkeys (*Chlorocebus aethiops sabeus*). Consistent with the human disease, T2D monkeys have increased plasma and CSF glucose levels as they transition from normoglycemia to pre-diabetic and diabetic states. Although plasma levels of acylcarnitines and amino acids remained largely unchanged, peripheral hyperglycemia correlated with decreased CSF acylcarnitines and CSF amino acids, including branched chain amino acid (BCAA) concentrations, suggesting profound changes in cerebral metabolism coincident with systemic glucose dysregulation. Moreover, CSF Aβ40 and CSF Aβ42 levels decreased in T2D monkeys, a phenomenon observed in the human course of AD which coincides with increased amyloid deposition within the brain. In agreement with our previous studies in mice, CSF Aβ40 and CSF Aβ42 were highly correlated with CSF glucose levels, suggesting that glucose levels in the brain are associated with changes in Aβ metabolism. Interestingly, CSF Aβ40 and CSF Aβ42 levels were also highly correlated with plasma but not CSF lactate levels, suggesting that plasma lactate might serve as a potential biomarker of disease progression in AD. Moreover, CSF glucose and plasma lactate levels were correlated with CSF amino acid and acylcarnitine levels, demonstrating alterations in cerebral metabolism occurring with the onset of T2D. Together, these data suggest that peripheral metabolic changes associated with the development of T2D produce alterations in brain metabolism that lead to early changes in the amyloid cascade, similar to those observed in pre-symptomatic AD.

## Introduction

Rates of type 2 diabetes (T2D) and Alzheimer’s disease (AD) are reaching epidemic proportions and are expected to continue to rise over the next several decades [1]. T2D is a metabolic disorder characterized by elevated fasting plasma glucose levels, increased insulin levels, insulin resistance, and beta cell dysfunction with the majority of changes occurring 5-10 years before clinical diagnosis [2]. Similarly, pathological hallmarks of AD, including the extracellular aggregation of amyloid β (Aβ) into amyloid plaques and the intracellular accumulation of the tau protein into neurofibrillary tangles (NFTs), begin decades before cognitive decline and clinical diagnosis [3–5]. While both are considered diseases of aging and mechanisms linking the two conditions remain elusive, epidemiological and cross-sectional studies suggest that individuals with T2D have a 2 – 4 fold increased risk for developing AD and dementia and show increased AD pathology [6–8]. Preclinical studies in mouse models of cerebral amyloidosis suggest that systemic hyperglycemia increases Aβ levels within the hippocampal interstitial fluid (ISF) by 25%; an effect that is amplified when plaques are already present in the brain during the hyperglycemia challenge [9, 10]. Moreover, mouse plasma glucose, ISF glucose, and ISF Aβ are highly correlated, and elevated glucose levels drive Aβ production in the hippocampus, in an activity dependent manner [9]. Conversely, systemic hyperinsulinemia at post-prandial or supra-physiological levels, only modestly increase ISF Aβ levels. This suggests that changes in glucose, rather than insulin, correlates more closely with brain Aβ levels [10]. Although these studies suggest a mechanistic link between T2D and AD, rodent models of AD do not fully recapitulate the human course of disease, and it is important to translate these observations to primates. Similar to humans, many non-human primate species develop T2D and AD pathology with age [11, 12], and thus represent an important translational tool for examining the metabolic relationship between the two conditions.

Branched chain amino acids (BCAAs), including leucine, isoleucine, and valine, are essential amino acids necessary for protein synthesis, but when found in excess, impact energy homeostasis [13–15]. Recent work demonstrated that elevated dietary BCAA intake is associated with obesity and insulin resistance in both humans and rodents [16, 17], and plasma BCAA levels are highly predictive of T2D development in normoglycemic individuals [18]. Elevated levels of circulating BCAAs are associated with suppressed mitochondrial β-oxidation, reduced glucose tolerance, increased insulin resistance, and increased de novo lipogenesis, making BCAAs a potential biomarker of metabolic disease [17, 19]. BCAAs are also integral to healthy brain function, due to their roles in neurotransmitter biosynthesis, protein synthesis, and energy production. Alterations in BCAA levels in plasma and CSF have been implicated in AD pathology, with conflicting evidence on whether they are helpful or harmful to disease progression [20]. Nevertheless, alterations in energy homeostasis and BCAA catabolism represent one potential link between T2D and AD.

Acylcarnitines are byproducts of mitochondrial fatty acid, amino acid and glucose catabolism that serve as useful biomarkers of metabolic changes [21]. Changes in the plasma acylcarnitine profile have been observed in obesity, T2D, and insulin resistance, representing alterations in several metabolic pathways [21, 22]. Moreover, acylcarnitines are key energy substrates in the brain, especially during fasting conditions when glucose levels are low [21]. In AD patients, plasma levels of acylcarnitines are decreased, suggesting perturbations in energy metabolism that may be central to AD pathogenesis [23].

Here, we applied comprehensive metabolic profiling tools to T2D, PreD, and healthy control (Ctrl) monkeys to explore the relationship between T2D and AD pathology. We analyzed plasma and CSF amino acids and acylcarnitine concentrations and explored how these changes related to CSF Aβ40 and Aβ42 levels, which are established biomarkers of disease in AD.

## Methods

### Animals

The monkeys used in this study were sourced from a multigenerational pedigreed colony of vervet monkeys (*Chlorocebus aethiops sabeus*; age = 16.5 – 23.5 years old). Veterinary and research staff categorized the monkeys as either healthy (Ctrl; n=4), pre-diabetic (PreD; n=4), or type-2 diabetic (T2D; n=5) according to repeated fasting glucose measurements and American Diabetes Association criteria [2] and were selected to be matched by age, bodyweight, and adiposity as measured by waist circumference. PreD and T2D categorization was only made after ≥2 consecutive fasting glucose values were ≥100 mg/dL or ≥126 mg/dL respectively. T2D monkeys were maintained with insulin therapy, and all T2D monkeys in study had been diagnosed and treated for a minimum of 2 years. Monkeys were fed a commercial laboratory primate chow diet (Laboratory Diet 5038; LabDiet, St. Louis, MO), with daily supplemental fresh fruits and vegetables. This standard laboratory diet is comprised of 13% calories from fat; 69% calories from carbohydrates; and 18% of calories from protein. All samples were collected after 16 hours fasting and withdrawal from all exogenous insulin. Cerebrospinal fluid (CSF) was collected via puncture of the atlanto-occipital space, and plasma samples were collected from the femoral vein.

All animal procedures were performed on a protocol approved by the Wake Forest University Institutional Animal Care and Use Committee according to recommendations in the Guide for Care and Use of Laboratory Animals (Institute for Laboratory Animal Research) and in compliance with the USDA animal Welfare Act and Animal Welfare Regulations (Animal Welfare Act as Amended; Animal Welfare Regulations).

### AD Biomarkers

Aβ_40_ and Aβ_42_ levels from CSF samples were assayed using sandwich ELISAs as previously described [24, 25]. Briefly, Aβ_40_ & Aβ_42_ were quantified using monoclonal capture antibodies targeted against amino acids 45-50 (HJ2) or 37-42 (HJ7.4), respectively. For detection, both Aβ40 and Aβ42 used a biotinylated monoclonal antibody against the central domain (HJ5.1B), followed by incubation with streptavidin-poly-HRP-40. Assays were developed using Super Slow TMB (Sigma) and the plates read on a Bio-Tek Synergy 2 plate reader at 650nm.

### Metabolomics, Lipids, Glucose, and Lactate Measures

Glucose and lactate measurements within the plasma and CSF were quantified using a YSI 2900 analyzer as previously described [9]. A detailed description of blood and CSF sample preparation and coefficients of variation for these assays has been published [16, 26]. Insulin was measured by ELISA (Mercodia, Uppsala, Sweden) in plasma and CSF samples. Total cholesterol, high density lipoprotein (HDL) and low-density lipoprotein (LDL) cholesterol, and triglycerides were measured with kits from Roche Diagnostics (Indianapolis, IN) and free fatty acids (total) and ketones (total and 3-hydroxybutyrate) with kits from Wako (Richmond, VA). ApoB associated cholesterol was calculated as the total cholesterol minus HDLC. Plasma and CSF acylcarnitines and amino acids were analyzed by MS/MS as described previously [27–31].

### Data Analysis

Data were analyzed using one-way ANOVA and correlations were determined by Pearson’s correlation coefficient, *r*. To determine the relative relationship between CSF Glucose, CSF Aβ_42_, and plasma lactate and CSF analytes, we transformed each data point to represent its value relative to the control group mean [% control mean value = 100*(x/control mean), where x = any given data point]. Data are represented as means ± SEM. Tukey’s post hoc tests were used when appropriate.

## Results

### Metabolic profile of normoglycemic, pre-diabetic, and T2D monkeys

Monkeys were older, ranging from 16-23 years (Table 1), which is typical age of onset for metabolic diseases. There were no differences in body weight or waist circumference between groups (Table 1). PreD and T2D monkeys had elevated fasting blood glucose levels compared to normoglycemic controls (Table 1; p<0.0001, F(2, 9)=39.17), but there were no differences in fasting insulin levels (Table 1). While HOMA scores were elevated in PreD and T2D monkeys, the differences were not significant (Table 1). Additionally, lipid measures illustrated higher triglycerides in T2D monkeys compared to PreD or Ctrl. Together, elevated fasting blood glucose was the most notable finding delineating the Ctrl, PreD, and T2D cohorts.

**Table 1.** Demographic and metabolic characteristics of monkeys included in study.

|  | Ctrl Mean (SEM) | PreD Mean (SEM) | T2D Mean (SEM) | One-Way ANOVA |
| --- | --- | --- | --- | --- |
| Age (years) | 20.25 (±1.109) | 18.75 (±0.6292) | 18.00 (±1.304) | p = 0.3800 |
| Body weight (kg) | 5.553 (±0.5525) | 6.618 (±0.9210) | 5.414 (±0.4937) | p = 0.4100 |
| Waist circumference (cm) | 36.98 (±1.719) | 41.94 (±3.861) | 38.08 (±3.441) | p = 0.5283 |
| **Fasting glucose (mg/dL)** | **67.84 (±5.097)** | **113.80 (±4.308)** | **161.10 (±11.05)** | **p < 0.0001** |
| Fasting insulin (μIU/mL) | 20.83 (±4.806) | 37.43 (±8.854) | 26.63 (±8.191) | p = 0.3284 |
| HOMA Score (AU) | 3.613 (±1.048) | 10.61 (±2.721) | 10.64 (±3.525) | p = 0.1512 |
| TPC (mg/dL) | 171.3 (±10.06) | 198.9 (±33.75) | 164.3 (±9.067) | p = 0.4945 |
| **Triglyceride (mg/dL)** | **74.33 (±2.541)** | **59.75 (±7.417)** | **103.80 (±16.19)** | **p = 0.0407** |
| HDLC (mg/dL) | 60.59 (±4.389) | 65.50 (±5.939) | 59.83 (±3.657) | p = 0.6702 |
| ApoB-associated cholesterol (mg/dL) | 110.70 (±6.232) | 95.22 (±8.152) | 104.40 (±6.809) | p = 0.3633 |
| TPC/HDLC | 2.845 (±0.093) | 2.360 (±0.130) | 2.785 (±0.148) | p = 0.0646 |

Ctrl = Control; PreD = Pre-Diabetic; T2D = Type-2 Diabetic; HOMA = Homeostatic model assessment; TPC = Total plasma cholesterol; HDLC = High-density lipoprotein cholesterol

### Increased CSF glucose levels correlated with decreased CSF Aβ_40_ and Aβ_42_ concentrations in T2D monkeys

T2D monkeys have elevated plasma glucose (Figure 1A; p=0.0036, F(2, 10)=10.39) and CSF glucose concentrations (Figure 1B; p<0.0001, F(2, 9)=32.31) compared to normoglycemic controls, while PreD had intermediate values. Data analysis revealed that plasma and CSF glucose levels had a strong positive correlation (Figure 1C; p<0.0002, r=0.8739, R^2^=0.7637), which is consistent with observations from our preclinical rodent models [9]. We also observed that plasma lactate levels were lower in T2D monkeys, however no differences in CSF lactate concentrations were observed (Figure 1D and E; p=0.0395, F(2, 10)=4.544). We previously demonstrated that hyperglycemic APP/PS1 mice, a model of cerebral amyloidosis, have elevated Aβ within the brain’s ISF [9, 10], and nonhuman primates with T2D have increased Aβ deposition in several brain regions [32]. T2D monkeys have decreased CSF Aβ_40_ (Figure 1F; p=0.0280, F(2, 9)=5.030;) and CSF Aβ_42_ concentrations (Figure 1J; p=0.0342, F(2, 9)=5.463). Interestingly, both CSF Aβ_40_ and CSF Aβ_42_ were highly correlated with CSF glucose (Figure 1G; p=0.0400, r=-0.5979, R^2^=0.3575 and Figure 1K; p=0.0489, r=-0.5782, R^2^=0.3343 respectively) and plasma lactate (Figure 1H; p=0.0027, r=0.7820, R^2^=0.6115 and Figure 1L p=0.0013, r=0.8133, R^2^=0.6614; respectively) but not with plasma insulin (Figure 1I & 1M). CSF insulin levels were undetectable. Given that decreased CSF Aβ is indicative of increased plaque formation within the brain [33], these data indicate that T2D monkeys display biomarkers of early amyloid deposition and pre-symptomatic AD triggered by a state of energy dysregulation, which is consistent with previous work in rodent models [9, 10].

**Figure 1.**
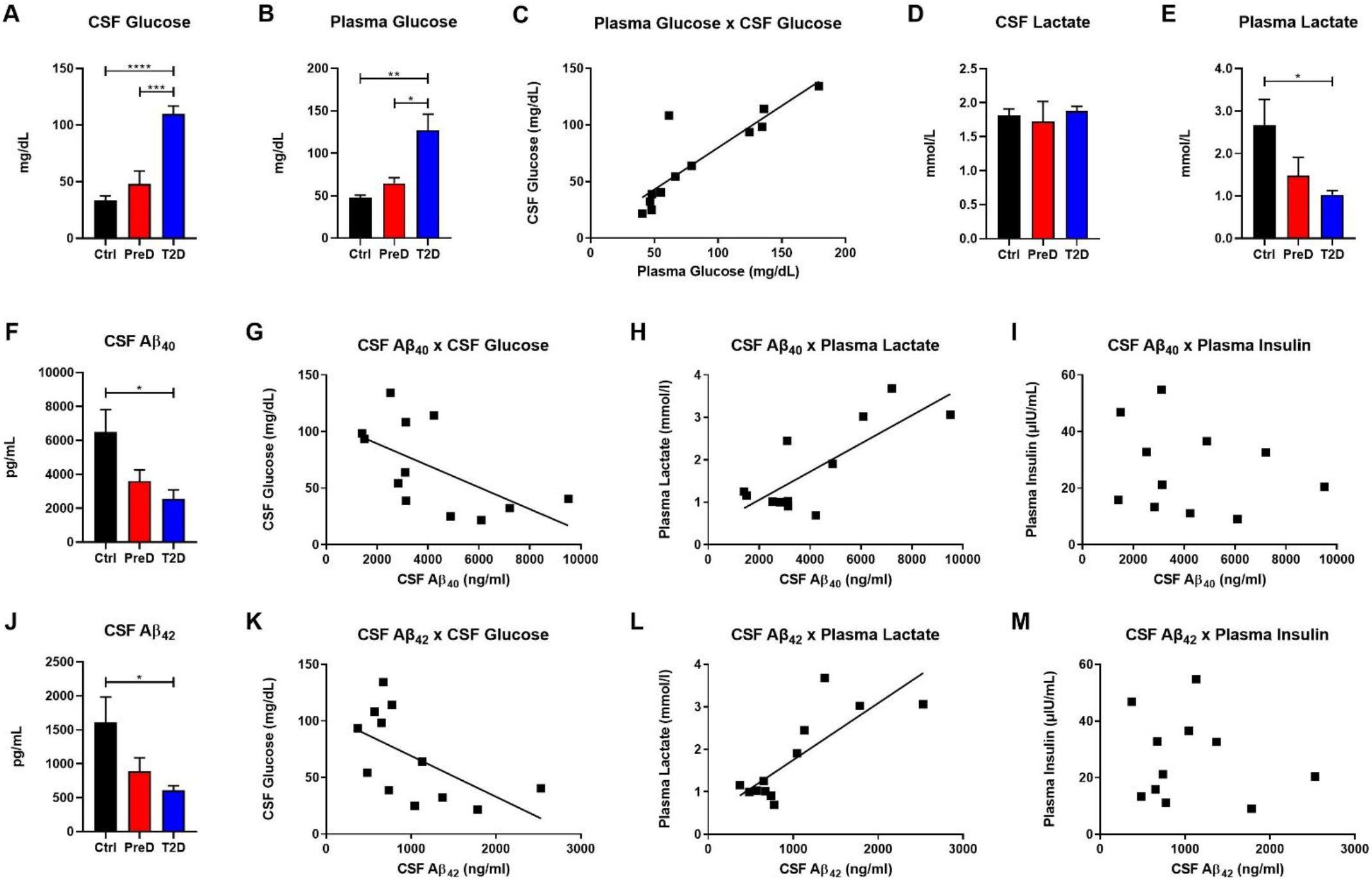
T2D monkeys have significantly decreased CSF Aβ_40_ and CSF Aβ_42_ concentrations, which is correlated with CSF glucose and plasma lactate concentrations. **(A)** Plasma glucose and **(B)** CSF glucose concentrations were significantly elevated in T2D monkeys. **(C)** Plasma and CSF glucose concentrations were significantly correlated. **(D)** T2D monkeys had significantly decreased plasma lactate, but not **(E)** CSF lactate. **(F)** T2D monkeys had significantly decreased Aβ_40_, which was correlated with **(G)** CSF glucose and **(H)** plasma lactate, but not **(I)** plasma insulin concentrations. **(J)** Similarly, T2D monkeys had significantly decreased Aβ_42_ concentrations, which was correlated with **(K)** CSF glucose and **(L)** plasma lactate, but not **(M)** plasma insulin concentrations..* p < 0.05; ** p < 0.01; *** p < 0.001; **** p < 0.0001; one-way ANOVA with Tukey’s *post hoc* test. Values represent mean ± SEM; solid lines represent statistically significant correlations (p<0.05).

### T2D and PreD monkey show decreased amino acid and acylcarnitines levels in the CSF but not the plasma

Because plasma amino acid and acylcarnitine levels are linked to metabolic dysfunction in T2D [22, 34], the levels of amino acids (AA) in both the plasma and CSF were quantified to further explore the energy imbalance associated with T2D. In examining the AA concentrations by their functional groups, PreD and T2D monkeys had lower concentrations of amino acids in the CSF. Amino acids were further stratified into branched-chain AA (BCAAs, Figure 2A; p=0.0038, F(2, 9)=11.04;), total AA (Figure 2B; p=0.0081, F(2, 9)=8.638), essential AA (Figure 2C; p=0.0065, F(2, 9)=9.280), aromatic AA (Figure 2D; p=0.0046, F(2, 9)=10.36), and basic AA (Figure 2E; p=0.0087, F(2, 9)=8.421). In the CSF, all AA groups were lower in T2D, with the exception of acidic AA where no changes were detected (Figure 2F). Interesting, no differences in plasma AA concentrations were detected in any of the AA categories (Figure 2GL). Interestingly, when individual AAs were measured in plasma, there was a trend towards an increase in levels of the BCAAs valine and leucine/isoleucine (Supplemental Figure 2F; p=0.0977 and Supplemental Figure 2M; p=0.1675, respectively), suggesting peripheral metabolic perturbations were present, although the differences in CSF valine and leucine/isoleucine were more striking (Supplemental Figure 1F; p=0.0219 and Supplemental Figure 1M; p=0.0524).

**Figure 2.**
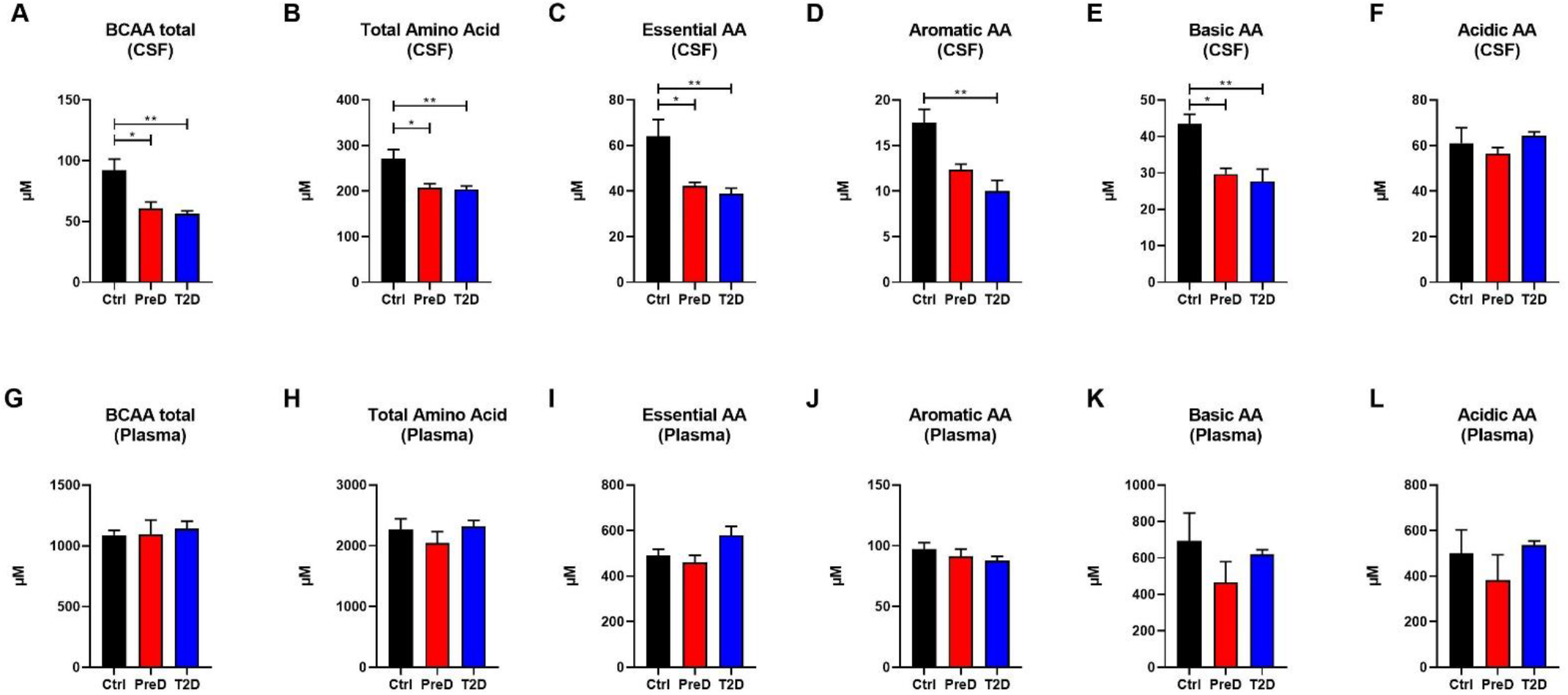
AA concentrations are decreased in the CSF but not plasma of T2D monkeys. **(A)** T2D monkeys had significantly decreased BCAA, **(B)** total amino acid concentrations, **(C)** essential AAs, **(D)** aromatic AAs, and **(E)** basic AAs, but not **(F)** acidic AAs. **(G-H)** Conversely, plasma amino acid concentrations were not different between groups.* p < 0.05; ** p < 0.01; *** p < 0.001; **** p < 0.0001; one-way ANOVA with Tukey’s *post hoc* test. Values represent mean ± SEM.

Acylcarnitines are derived from the mitochondrial oxidation of fatty acids, carbohydrates, and amino acids [22]. Several studies have shown that T2D patients have elevated plasma acylcarnitine concentrations compared to healthy controls[35]. Here, T2D monkeys had lower total acylcarnitine concentrations in the CSF (Figure 3A; p=0.0208, F(2, 9)=6.139;) but no differences in plasma total acylcarnitine levels (Figure 3C), a pattern consistent with the AA data. In the CSF, T2D monkeys also had lower short-chain acylcarnitine concentrations (Figure 3B; p=0.0245, F(2, 9)=5.764). However due to variability in the control monkeys, medium- and long-chain acylcarnitine concentrations were consistently lower in T2D monkeys, but the difference did not reach significance [35] (Figure 3C-D). Again, plasma concentrations remained comparable between groups (Figure 3F-H). Together, this data suggests that fuel metabolism is altered in the brains of T2D monkeys compared to normoglycemic controls.

**Figure 3.**
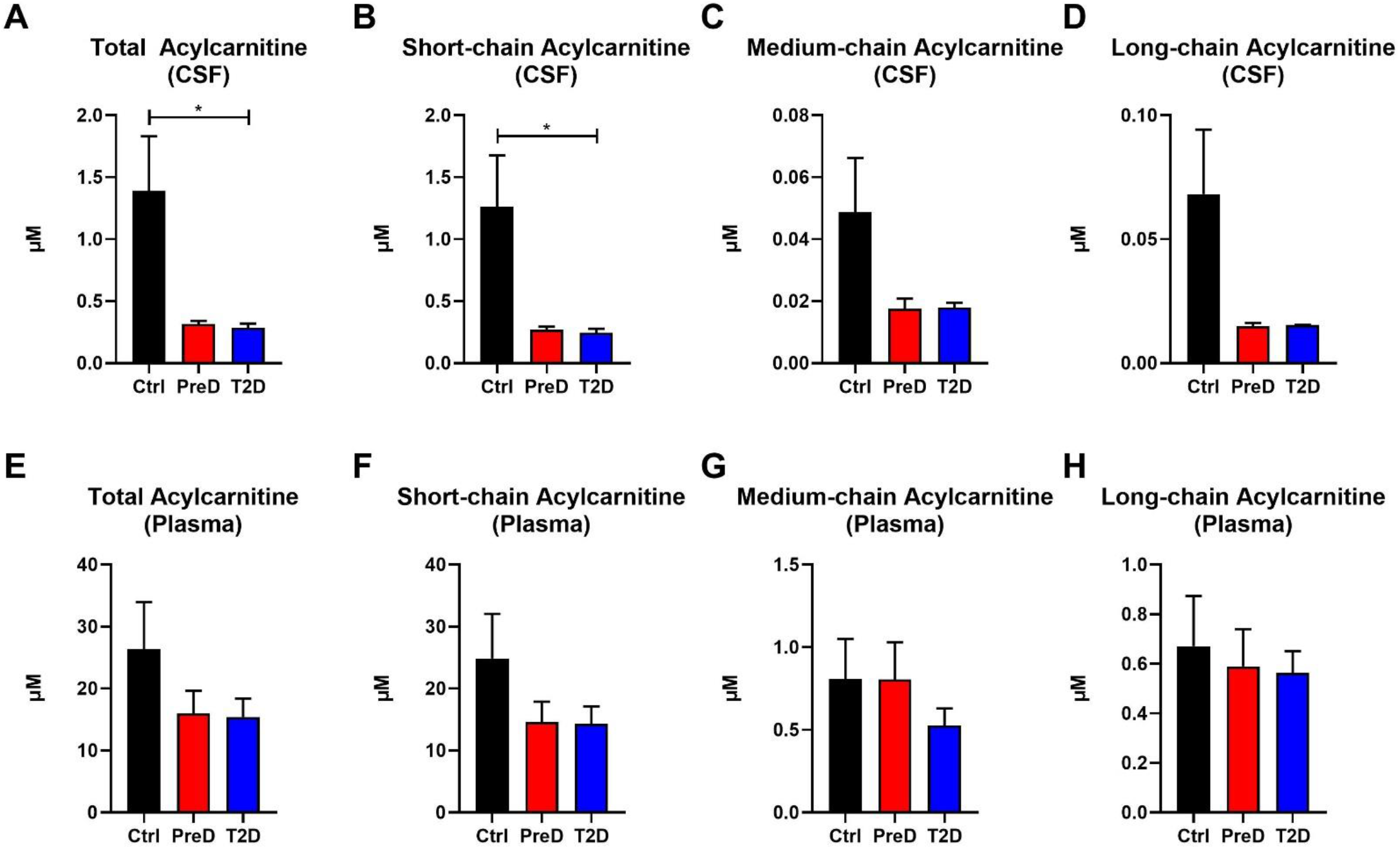
Acylcarnitine concentrations are decreased in the CSF but not plasma of T2D monkeys. **(A-B)** Total and short-chain acylcarnitine concentrations were significantly decreased in the plasma of T2D and IR monkeys, **(C-D)** however medium- and long-chain acylcarnitines were not significantly different. **(E-H)** There were no differences in plasma acylcarnitine concentrations between groups.* p < 0.05; one-way ANOVA with Tukey’s *post hoc* test. Values represent mean ± SEM.

### Metabolic dysregulation in the brain is associated with changes in CSF glucose, CSF Aβ_42_, and plasma lactate

Lastly, we investigated the relationship between differences in CSF amino acids and CSF acylcarnitines as a function of CSF glucose, CSF Aβ_42_, and plasma lactate concentrations (Figure 4) to further elucidate the interaction between early metabolic changes in T2D with early biomarker alterations in AD. There was an overall negative relationship between CSF glucose and Aβ_40_, Aβ_42_, total AA, essential AA, BCAA, aromatic AA, basic AA, short-chain acylcarnitine, and total acylcarnitine (Figure 4A). Thus, as CSF glucose increases as observed in PreD and T2D, concentrations of amino acids, acylcarnitines, and Aβ all decrease in the CSF. Next, CSF Aβ_42_ was correlated with CSF Aβ_40_ and CSF BCAA (Figure 4B), reinforcing the relationship between BCAAs and AD pathology. Lastly, plasma lactate concentrations correlated with CSF Aβ_40_ and Aβ_42_, CSF essential AA and BCAA, and CSF short-chain and total acylcarnitine concentrations (Figure 4C), demonstrating plasma lactate might be a potential biomarker for early changes in T2D and pre-symptomatic AD. Taken together, alterations in cerebral metabolism co-vary with changes in plasma glucose, plasma lactate, and CSF Aβ_42_ which highlight the importance of metabolic changes in the pathogenesis of T2D and AD.

**Figure 4.**
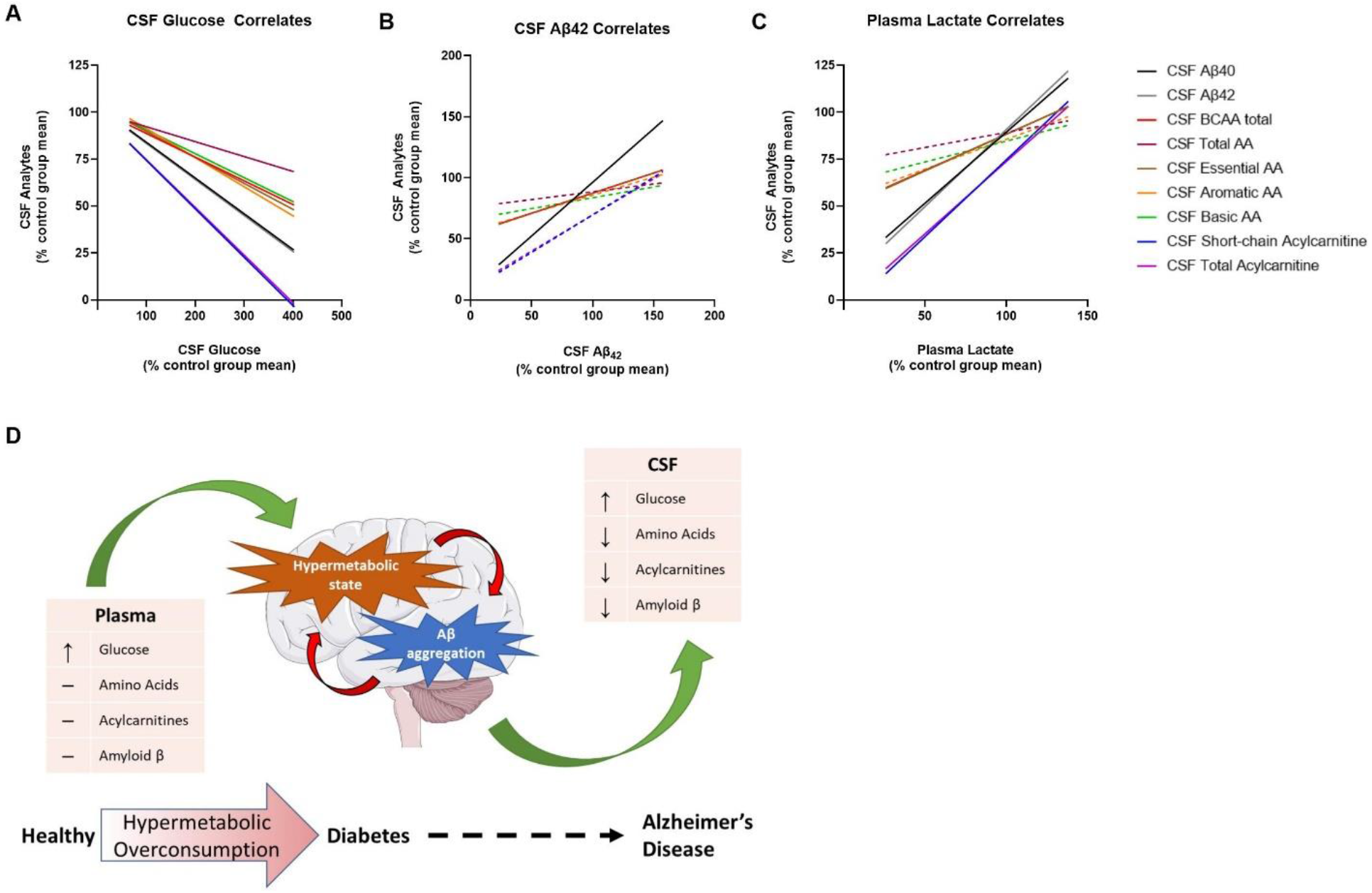
Metabolic dysregulation is associated with changes in plasma lactate, CSF glucose, and CSF Aβ_40_ and Aβ_42_. Here, each data point has been converted into a value that represents the % of the control group’s mean [value = 100*(x/control group mean)]. **(A)** Normalized CSF glucose values were negatively correlated with several normalized CSF analytes. **(B)** Normalized CSF Aβ_42_ was correlated with normalized CSF analytes. Lines represent nonlinear regression best fit curve; solid lines indicate statistical significance. **(C)** Normalized plasma lactate values were positively correlated with several normalized CSF analytes. **(D)** Thus, the model we propose here is that T2D moves the brain into a state of hypermetabolic overconsumption wherein the brain consumes increased energy, leading to lower concentrations of Aβ_42_ in the CSF, which is indicative of increased Aβ aggregation. Solid lines regression lines denote statistical significance.

## Discussion

In this study, elevated fasting blood glucose levels associated with the onset of T2D elicit changes in brain metabolism and correlate with changes in the amyloid cascade, an early indicator of presymptomatic AD [8, 9]. T2D monkeys had lower CSF Aβ40 and Aβ42 levels, which is indicative of increased amyloid deposition within the brain [33]. In agreement with previous rodent studies [9, 10], CSF Aβ_40_ and Aβ_42_ were highly correlated with CSF glucose levels, which suggests that increased glucose may be driving Aβ production and aggregation. Interestingly, CSF Aβ_40_ and Aβ_42_ levels also highly correlated with plasma lactate levels, which is consistent with published studies that show decreased plasma lactate correlates with AD severity [36, 37]. Moreover, T2D vervet monkeys had lower CSF acylcarnitine and CSF amino acids, while plasma levels were largely unchanged, suggesting either early changes in cerebral metabolism with the onset of T2D or changes in cerebral delivery/transport of certain metabolic fuels. Reduced CSF Aβ_40_ and Aβ_42_ levels in T2D monkeys correlated with higher plasma and CSF glucose concentrations, signifying onset of pre-symptomatic AD. Lastly, we showed that CSF amino acids and acylcarnitines were negatively correlated with CSF glucose and positively correlated with CSF Aβ_40_, CSF Aβ_42_, and plasma lactate. Together, these data suggest that peripheral metabolic changes associated with diabetogenesis co-occur with alterations in brain metabolism. Moreover, these metabolic changes are associated with activation of the amyloid cascade typically associated with pre-symptomatic AD.

Our data further supports existing evidence that chronic hyperglycemia and metabolic dysfunction are a pathological link between T2D and AD. In humans, hyperglycemia increases dementia risk in both patients with and without diabetes, causes rapid progression from mild cognitive impairment (MCI) to symptomatic AD, and increases the rate of amyloid accumulation in the brain [8, 38]. Moreover, hyperglycemia and increased HbA1c levels correlate with memory impairment, decreased functional connectivity, and increased neuronal loss, independent of T2D or AD diagnosis [39]. Data from T2D monkeys illustrates the same phenomenon; elevated blood glucose levels increase CSF glucose which correlates with changes in CSF Aβ levels, presumably due to the sequestration of Aβ into amyloid plaques in the brain [40]. Our findings in the T2D monkeys also uncovered an interesting relationship between glucose, lactate, and Aβ which supports previous findings from rodent studies. Preclinical studies in mouse models of cerebral amyloidosis demonstrate that synaptic release of Aβ occurs in an activity dependent manner, where high levels of synaptic activity increase Aβ secretion [40–43]. Increased synaptic activity not only drives ISF Aβ release but also the release of lactate into the extracellular space [25]. According to the astrocyte neuron lactate shuttle, lactate is a preferred energy source for neurons to sustain excitatory neurotransmission and levels of ISF lactate correlate with neuronal activity. Our previous work demonstrated that hyperglycemia not only increases ISF glucose and ISF Aβ but also ISF lactate, illustrating that increased metabolic activity is linked with increased synaptic activity and Aβ release. Since a direct measure of lactate production in the brain of T2D monkeys was unattainable in this study, we explored how plasma and CSF levels changed with peripheral hyperglycemia. Interestingly, plasma lactate, but not CSF lactate, correlated with changes in CSF glucose and Aβ. In accordance with our previous work, we hypothesize that increased glucose metabolism is increasing neuronal activity within the brain and driving both the production of Aβ and the consumption of pyruvate and lactate as fuel. Although the changes in plasma lactate levels could be due to alterations in peripheral metabolism in the T2D monkeys, we propose a different mechanism where increased lactate consumption in the brain signifies a hyperactive and hypermetabolic brain state present in both T2D and AD (Figure 4D). Since the concentration gradient for lactate favors transport from brain to blood [44], decreased plasma lactate levels could reflect increased neuronal activity, lactate consumption, and Aβ production in the brain, which also makes plasma lactate levels a potential serum biomarker for AD, T2D, or both. In humans, a small cohort study established that serum lactate levels decreased in symptomatic AD, yet the authors attributed this finding to alterations in muscle metabolism, not brain [37]. Therefore, additional studies need to be performed to elucidate the role of plasma lactate in T2D and AD.

In the current study, we demonstrated that T2D monkeys have lower CSF acylcarnitine and amino acid concentrations (Figures 2 & 3). Several studies demonstrated that circulating levels of amino acids are positively correlated with obesity, insulin resistance, metabolic dysfunction, and T2D in humans and rodents [16–18]. Although our data demonstrates a trend towards an increase in the BCAAs valine and isoleucine/leucine, no differences in plasma amino acid concentrations were detected in PreD or T2D monkeys. This may be explained by the fact that the T2D monkeys in this study were fed a well-controlled, well balanced diet that did not replicate the traditional nutritional overconsumption seen humans with metabolic syndrome and T2D. Also, although the T2D monkeys were hyperglycemic, their insulin levels were unchanged, suggesting that the hyperglycemia may arise via a mechanism independent of insulin resistance. Studies reporting elevated plasma BCAAs in humans have involved obese and insulin resistant subjects [16, 17]. Therefore, future studies should explore the relationship between plasma and CSF amino acid levels in a non-human primate model of dietary induced metabolic syndrome and T2D.

Another explanation for the difference in CSF AA levels could be that the T2D brain is overconsuming amino acids as fuel or rapidly increasing protein synthesis. While glucose is the primary source of energy for the brain, the brain can readily use fatty-acids as energy substrates; however, this typically occurs with decreased glucose availability, such as fasting or starvation [45]. This could lead the brain to a state of hypermetabolic overconsumption if both glucose and fatty acid metabolism were upregulated. Furthermore, because many of the amino acids consumed by the brain are necessary for neurotransmitter biosynthesis or neurotransmission itself [46], we propose that increased amino acid consumption increases both synaptic activity and metabolic activity, leading to elevated Aβ production, oxidative stress, and Aβ aggregation. Our current data cannot discern if the decrease in amino acids and acylcarnitines in the CSF is due to increased oxidation, or by another means, such as altered amino acid transport, therefore additional studies will need to further elucidate mechanisms underlying the changes in CSF metabolites.

In conclusion, the data presented here show that in the progression from healthy to PreD to T2D, the brain moves into a state of altered metabolism that results in an increase in glucose and lowering of amino acids and acylcarnitines in the CNS. This metabolic mileu increase Aβ production and accelerates Aβ aggregation, which reciprocally escalates the disease cascades in T2D and AD. These findings shed further light on the metabolic link between T2D and AD pathology and how T2D progression could lead to AD-related pathology and cognitive decline.

## Acknowledgements

We would like to acknowledge the following grants: 1K01AG050719 (SLM), R01AG061805 (SLM), NCDRC Pilot Award (SLM), Harold and Mary Eagle Fund for Alzheimer’s Research (SLM). The authors gratefully acknowledge use of the Wake Forest Nonhuman Primate Program, funded by the National Center for Advancing Translational Sciences and Office of the Director (KK): UL1TR001420 and P40-OD010965.

---

**Supplementary Figure 1.**
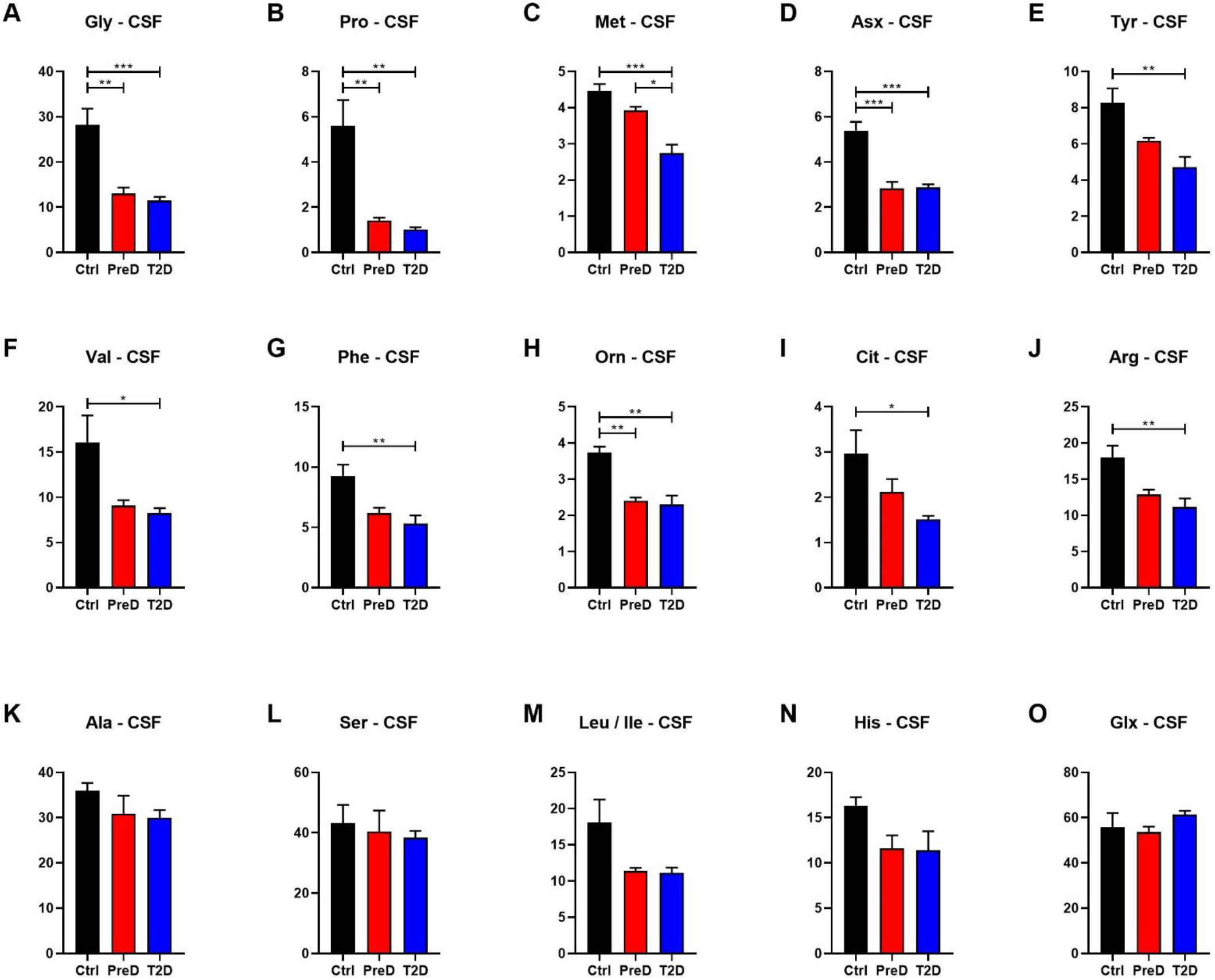
T2D monkeys had significantly decreased levels of several key essential, branched chain, and aromatic amino acids. T2D and PreD monkeys had significantly decreased **(A-E)** Gly, Pro, Met, Asx, and Tyr. **(F-J)** T2D monkeys had significantly decreased Val, Phe, Orn, Cit, and Arg. **(K-O)** There were no differences in Ala, Ser, Leu/Ile, His, or Glx between any groups.

**Supplemental Figure 2.**
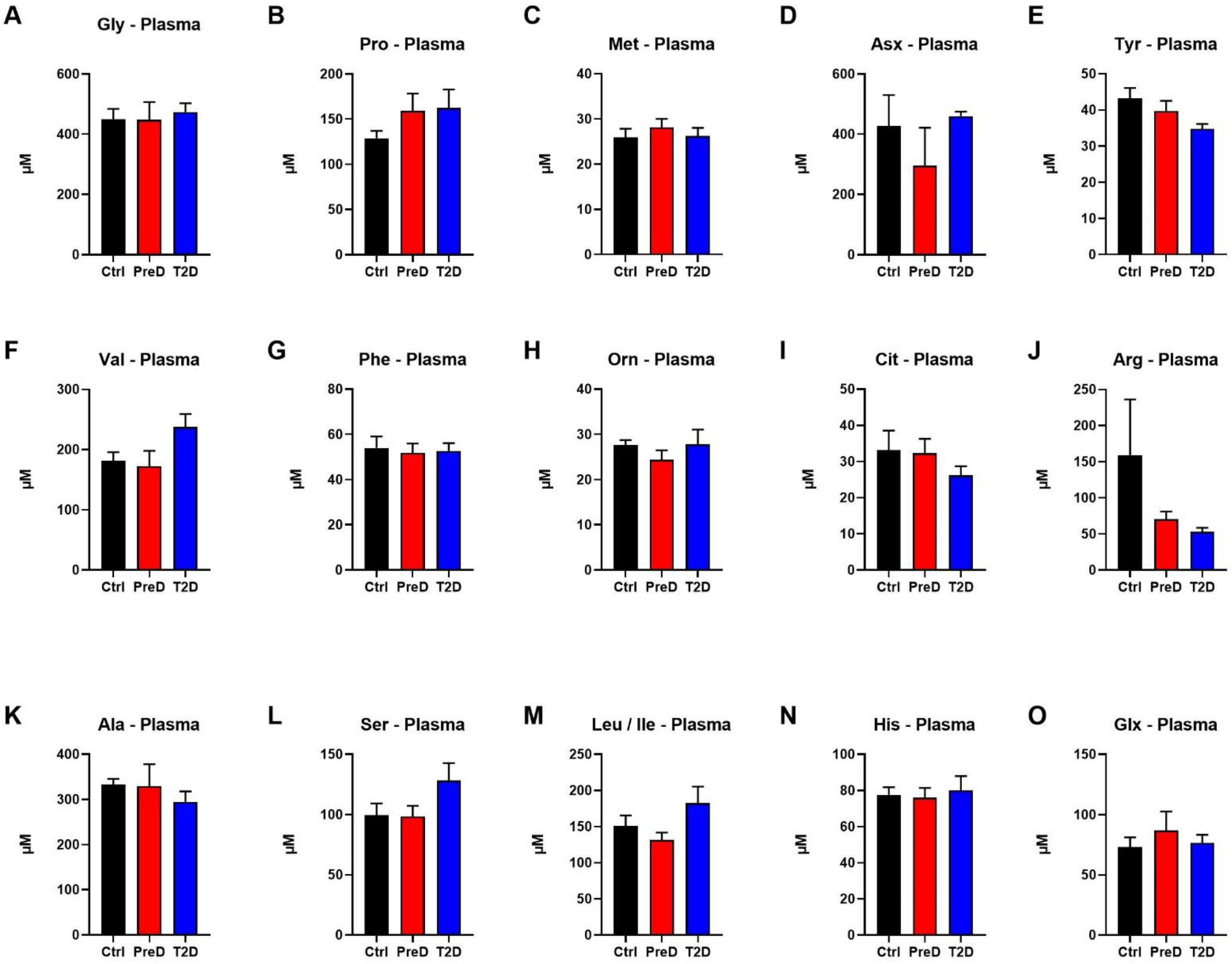
There were no differences in any single amino acid concentrations in the plasma between groups. **(A-O)** Amino acid concentrations were similar between Ctrl, PreD, and T2D monkeys.

**Supplemental Figure 3.**
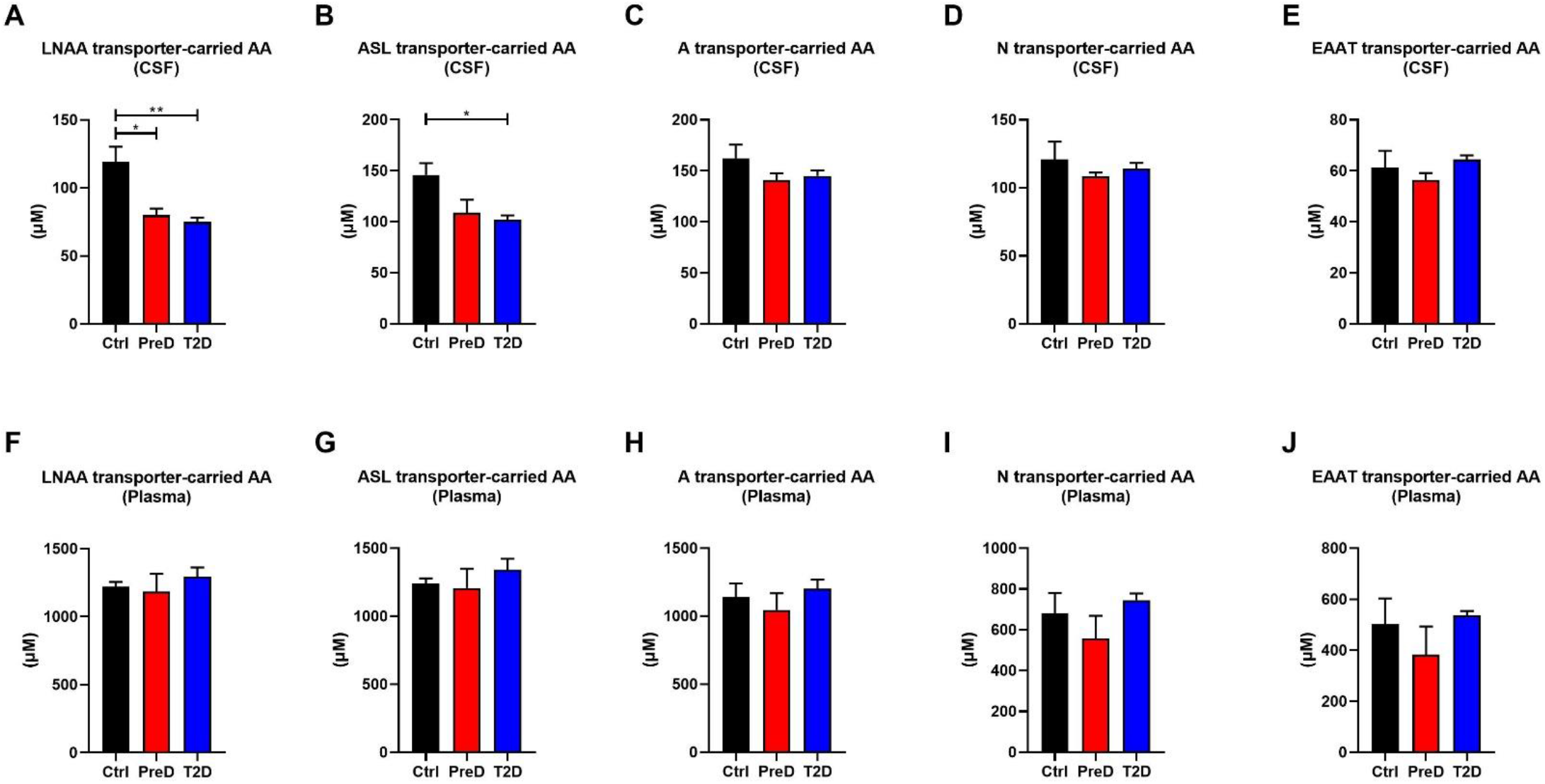
LNAA- and ASL-transported AAs are decreased in the CSF of T2D monkeys. **(A-B)** LNAA- and ASL-transported AAs are decreased in the CSF of T2D monkeys. **(C-D)** There were no differences in A-, N-, and EAAT-transported AA concentrations in CSF of PreD and T2D monkeys. **(F-J)** There were no differences in LNAA-, ASL-, A-, N-, and EAAT-transported AA concentrations in plasma of PreD or T2D monkeys

**Supplemental Table 1.** Levels of plasma and CSF amino acid data

|  | Ctrl Mean (SEM) | PreD Mean (SEM) | T2D Mean (SEM) | One Way ANOVA |
| --- | --- | --- | --- | --- |
| BCAA total (CSF) | 92.21 (±9.191) | 60.63 (±5.38) | 56.3 (±2.39) | 0.0038 |
| Total AA (CSF) | 271.3 (±19.48) | 208 (±7.834) | 203.7 (±7.605) | 0.0081 |
| Essential AA (CSF) | 64.19 (±7.218) | 42.21 (±1.656) | 38.85 (±2.435) | 0.0065 |
| Aromatic AA (CSF) | 17.54 (±1.449) | 12.35 (±0.6212) | 10.03 (±1.182) | 0.0046 |
| Basic AA (CSF) | 43.46 (±2.695) | 29.73 (±1.54) | 27.75 (±3.332) | 0.0087 |
| Acidic AA (CSF) | 61.19 (±6.614) | 56.41 (±2.754) | 64.34 (±1.685) | 0.4606 |
| Essential AA (Plasma) | 490.6 (±26.86) | 460.4 (±30.71) | 579.3 (±38.81) | 0.0708 |
| Total AA (Plasma) | 2267 (±176.2) | 2045 (±187.1) | 2318 (±97.31) | 0.4315 |
| BCAA total (Plasma) | 1089 (±37.34) | 1096 (±116.4) | 1140 (±63.76) | 0.8752 |
| Aromatic AA (Plasma) | 97.29 (±5.295) | 91.48 (±5.884) | 87.5 (±3.871) | 0.3972 |
| Basic AA (Plasma) | 693 (±154.2) | 466.6 (±112.8) | 621.2 (±24.27) | 0.3415 |
| Acidic AA (Plasma) | 501.6 (±101.5) | 382.7 (±111.1) | 536.8 (±17.37) | 0.4001 |

**Supplemental Table 2.**
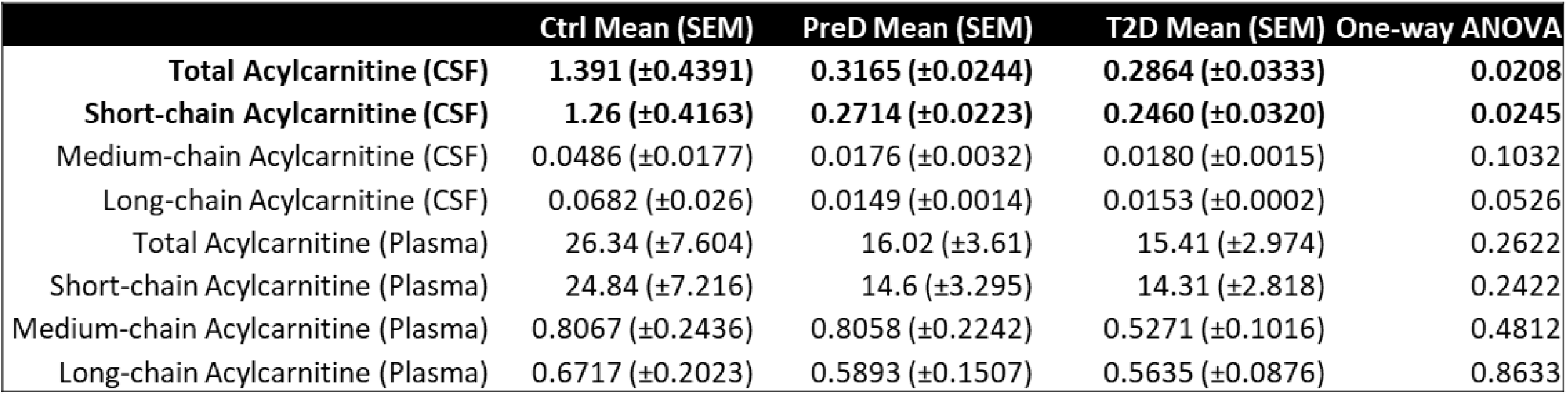
Levels of acylcarnitines in the CSF and plasma

**Supplemental Table 3.**
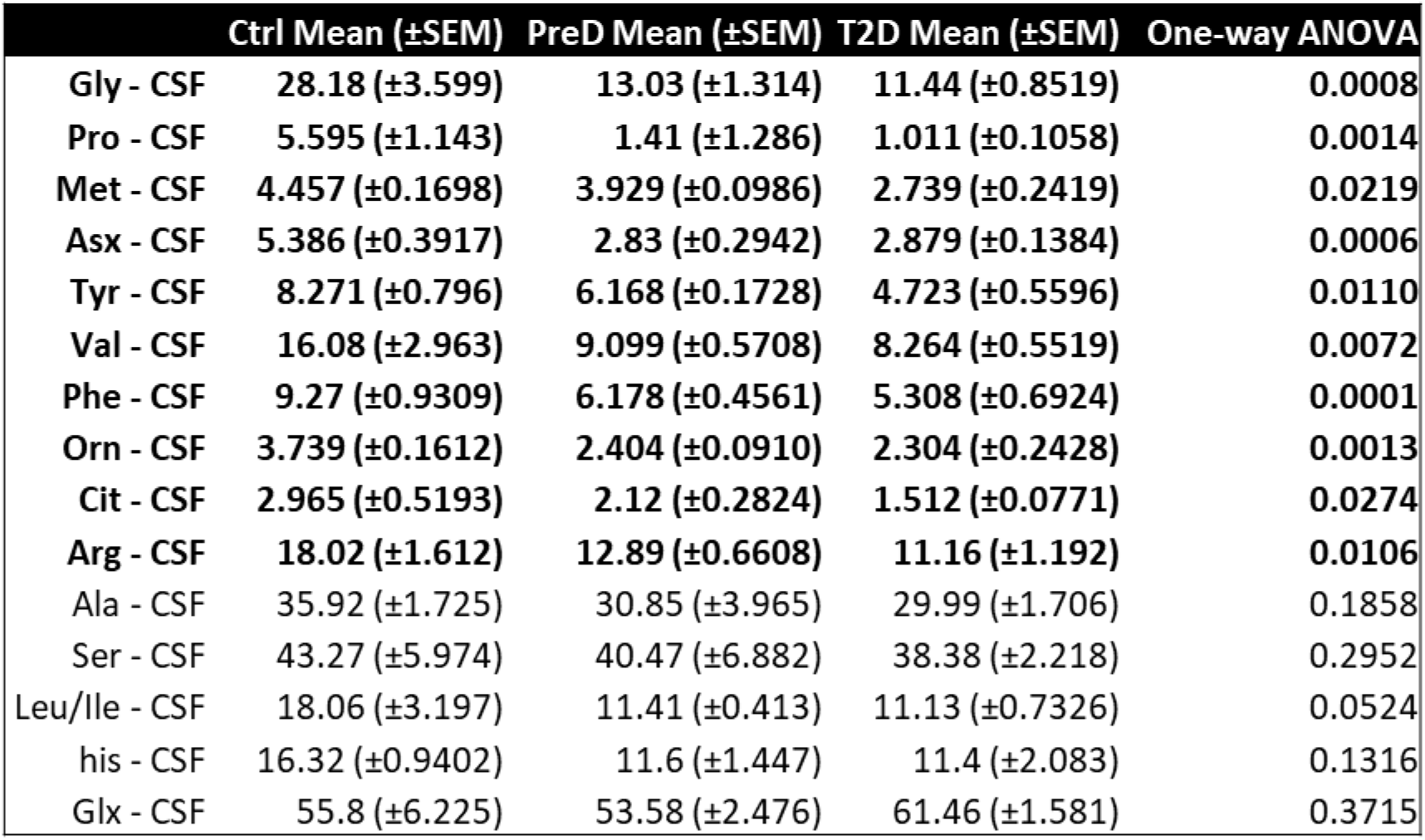
Levels of individual amino acids in the CSF.

| | Ctrl Mean (±SEM) | PreD Mean (±SEM) | T2D Mean (±SEM) | One-way ANOVA |
| --- | --- | --- | --- | --- |
| Gly - CSF | 28.18 (±3.599) | 13.03 (±1.314) | 11.44 (±0.8519) | 0.0008 |
| Pro - CSF | 5.595 (±1.143) | 1.41 (±1.286) | 1.011 (±0.1058) | 0.0014 |
| Met - CSF | 4.457 (±0.1698) | 3.929 (±0.0986) | 2.739 (±0.2419) | 0.0219 |
| Asx - CSF | 5.386 (±0.3917) | 2.83 (±0.2942) | 2.879 (±0.1384) | 0.0006 |
| Tyr - CSF | 8.271 (±0.796) | 6.168 (±0.1728) | 4.723 (±0.5596) | 0.0110 |
| Val - CSF | 16.08 (±2.963) | 9.099 (±0.5708) | 8.264 (±0.5519) | 0.0072 |
| Phe - CSF | 9.27 (±0.9309) | 6.178 (±0.4561) | 5.308 (±0.6924) | 0.0001 |
| Orn - CSF | 3.739 (±0.1612) | 2.404 (±0.0910) | 2.304 (±0.2428) | 0.0013 |
| Cit - CSF | 2.965 (±0.5193) | 2.12 (±0.2824) | 1.512 (±0.0771) | 0.0274 |
| Arg - CSF | 18.02 (±1.612) | 12.89 (±0.6608) | 11.16 (±1.192) | 0.0106 |
| Ala - CSF | 35.92 (±1.725) | 30.85 (±3.965) | 29.99 (±1.706) | 0.1858 |
| Ser - CSF | 43.27 (±5.974) | 40.47 (±6.882) | 38.38 (±2.218) | 0.2952 |
| Leu/Ile - CSF | 18.06 (±3.197) | 11.41 (±0.413) | 11.13 (±0.7326) | 0.0524 |
| his - CSF | 16.32 (±0.9402) | 11.6 (±1.447) | 11.4 (±2.083) | 0.1316 |
| Glx - CSF | 55.8 (±6.225) | 53.58 (±2.476) | 61.46 (±1.581) | 0.3715 |

**Supplemental Table 4.** Levels of individual amino acids in the plasma.

|  | Ctrl Mean (±SEM) | PreD Mean (±SEM) | T2D Mean (±SEM) | One-way ANOVA |
| --- | --- | --- | --- | --- |
| Gly - Plasma | 449.7 (±34.58) | 447.5 (±59.17) | 474 (±28.84) | 0.8744 |
| Ala - Plasma | 333.5 (±12.02) | 329.3 (±48.75) | 294.9 (±23.01) | 0.3960 |
| Ser - Plasma | 99.55 (±9.633) | 68.38 (±8.915) | 128.3 (±14.44) | 0.6486 |
| Pro - Plasma | 128.9 (±8.201) | 159.5 (±18.97) | 192.6 (±20.31) | 0.6178 |
| Val - Plasma | 181.7 (±13.95) | 172.6 (±25.2) | 237.6 (±21.61) | 0.1675 |
| Leu/Ile - Plasma | 151.4 (±14.04) | 131.8 (±10.05) | 182.6 (±22.68) | 0.5904 |
| Met - Plasma | 25.9 (±1.967) | 28.18 (±1.853) | 26.23 (±1.834) | 0.1701 |
| his - Plasma | 77.61 (±4.167) | 76.14 (±5.333) | 80.15 (±7.791) | 0.4078 |
| Phe - Plasma | 54.03 (±5.047) | 51.75 (±4.202) | 52.7 (±3.468) | 0.6806 |
| Tyr - Plasma | 43.26 (±2.859) | 39.73 (±2.795) | 34.8 (±1.359) | 0.0977 |
| Asx - Plasma | 428.4 (±101.8) | 295.8 (±125.8) | 460.2 (±14.43) | 0.3728 |
| Glx - Plasma | 73.2 (±7.929) | 86.96 (±15.68) | 76.64 (±6.648) | 0.2011 |
| Orn - Plasma | 27.66 (±1.076) | 24.41 (±2.059) | 27.81 (±2.259) | 0.9009 |
| Cit - Plasma | 33.09 (±5.468) | 32.32 (±3.948) | 26.17 (±2.527) | 0.9334 |
| Arg - Plasma | 159.3 (±76.89) | 70.26 (±10.74) | 53.02 (±5.703) | 0.0681 |

**Supplemental Table 5.** Levels of amino acids, grouped by AA transporter, in the plasma and CSF.

| | Ctrl Mean (±SEM) | PreD Mean (±SEM) | T2D Mean (±SEM) | One-way ANOVA |
| --- | --- | --- | --- | --- |
| **LNAA transporter AA (CSF)** | **119.0 (±11.3)** | **79.92 (±4.86)** | **74.96 (±3.264)** | **0.0034** |
| **ASL transporter AA (CSF)** | **146.0 (±11.46)** | **108.8 (±12.76)** | **101.9 (±4.35)** | **0.0137** |
| A transporter AA (CSF) | 162.3 (±13.34) | 140.7 (±6.61) | 145.1 (±5.184) | 0.2733 |
| N transporter AA (CSF) | 120.8 (±13.17) | 108.5 (±2.757) | 114.1 (±4.245) | 0.6340 |
| EAAT transporter AA (CSF) | 61.19 (±6.614) | 56.41 (±2.754) | 64.34 (±1.685) | 0.4606 |
| **LNAA transporter AA (Plasma)** | **1220 (±35.17)** | **1185 (±128.6)** | **1296 (±66.1)** | **0.6321** |
| **ASL transporter AA (Plasma)** | **1242 (±37.74)** | **1208 (±142.5)** | **1344 (±79.25)** | **0.5708** |
| A transporter AA (Plasma) | 1141 (±100.9) | 1046 (±123.5) | 1203 (±64.79) | 0.5176 |
| N transporter AA (Plasma) | 678.7 (±102.0) | 557.3 (±111.8) | 745.3 (±32.99) | 0.3055 |
| EAAT transporter AA (Plasma) | 501.6 (±101.5) | 382.7 (±111.1) | 536.8 (±17.37) | 0.4001 |

